# UCLA’s Competitive Edge Program Provides an Advantage to STEM Doctoral Students from Historically Excluded and Underrepresented Groups

**DOI:** 10.1101/2023.09.08.555984

**Authors:** Christina A. Del Carpio, Diana E. Azurdia

## Abstract

Universities benefit from recruiting and retaining diverse students, as it leads to more creative and rigorous problem solving. Efforts to improve the diversity, equity, and inclusion of graduate education have historically been focused on recruitment but are now shifting to retaining enrolled students. A low percentage of doctoral students, particularly those from historically underrepresented groups (URGs), complete their PhD within 10 years. To increase support of incoming doctoral students from URGs, UCLA established the Competitive Edge (CE) bridge program. CE provides six weeks of professional development and research training at the start of the PhD program. We surveyed 55+ first-year doctoral students (14 CE students, eight non-CE students from URGs, and 34 well-represented (WR) students) in STEM fields to understand CE’s effectiveness. We found that the CE program aided students in four areas that influence graduate student attrition: mentor/mentee relationship, socialization, finances, and preparedness. At the end of their first academic year, CE students reported that the program helped them in multiple areas relating to student success, such as mental wellbeing and sense of belonging. CE students reported a larger mean growth in seven of eight skills needed in graduate school compared to NonCE URG and WR students. Short answer responses revealed that NonCE students wished for more support in areas covered by the CE program, such as managing advising relationships and protecting mental health. Additionally, CE students received significant funding during the program. The CE program’s successful model at UCLA can be adapted to improve support for underrepresented doctoral students at other institutions.

## Introduction

Diversity among researchers is known to lead to a diversity of ideas and approaches that in turn leads to more creative and rigorous problem solving. Thus, it is in universities’ best interests to recruit and retain diverse students. However, historic efforts in graduate education have been more focused on recruitment and only recently began to shift towards support of matriculated students. In the literature on graduate student education, there are multiple studies focused on the reasons students leave graduate school (*e.g.* Bowlin, Sweat, Watts, & Throne, 2017; Hardré, Liao, Dorri, & Beeson Stoesz, 2019; Maher, Wofford, Roksa, & Feldon, 2017; Meara, Griffin, & Robinson, 2017; Rockinson-Szapkiw, 2019), but the literature is more limited on how institutions can counteract those factors to retain diverse talent. Some literature that focused on undergraduates explored the role of “bridge programs” to facilitate students in successfully applying to graduate schools (*e.g.* Mccoy, Winkle-wagner, & Winkle-wagner, 2020; Peteet et al., 2016), yet there are limited peer-reviewed publications on doctoral bridge programs that target admitted students. To meet the need for supporting incoming doctoral students from historically excluded and underrepresented groups (URGs), the University of California, Los Angeles (UCLA) has instituted the Competitive Edge (CE) bridge program. The program takes place over six weeks during the summer before typical doctoral programs begin. During this time, students receive professional and “soft” skills development, as well as research training. Additionally, CE gives students a chance to acclimate to graduate school and build community before the start of their Ph.D. program. In this article, we have surveyed 55+ first-year doctoral students in science, technology, engineering, and math (STEM) fields to gain insight into the effectiveness of CE as an approach to support diverse students in their transition to doctoral study.

### Why students leave graduate school

Part of the urgency to increase support and retention of students comes from data showing that, after 10 years from starting their graduate program, less than 60% of all doctoral students in all fields complete their PhD (Sowell, Zhang, Redd, & King, 2008). Additional data shows seven year completion rates to be similarly low for Hispanic/Latino students, and the rates for Black/African-American students to be as low as ∼50% (Sowell, Allum, & Okahana, 2015). It is also worth noting that other ethnic and racial minorities were not analyzed by Sowell et al. (2015) because there were too few students in the doctoral student body to reach a significant sample size. Increasing retention of historically underrepresented groups is also an important piece in actively creating a balance of people from diverse backgrounds at all levels of the academia hierarchy.

A meta analysis of 79 graduate student attrition studies identified four key themes that repeatedly emerge as reasons students leave: **1)** advisor-advisee relationship, **2)** socialization, **3)** finances, and **4)** preparedness (Bowlin et al., 2017).

#### Advisor-Advisee Relationship

The impact of the advising relationship on student retention is unsurprising given its central role in the model of graduate school. For example, in most STEM fields, students work within a given lab under the supervision of a principal investigator (PI), particularly so in Biology. (Note the terms PI, mentor, and advisor are typically used interchangeably in STEM PhD programs.) The advisor is charged with overseeing the student’s academic and professional growth, but also has a significant role in a student’s degree progress and future career of the student through dissertation project assignments, funding, and letters of recommendation. Given the central role of an advisor to a STEM PhD student’s work, it follows that having a constructive and supportive advisor-advisee relationship is crucial to a student’s experience and chances for success (Weiss, 1981, Girves & Wemmerus, 1988; Lovitts, 2001; Ruud, Saclarides, George-Jackson, & Lubienski, 2018; National Academies of Sciences, Engineering, and Medicine, 2019).

#### Socialization

Student experiences are also impacted by socialization with peers, faculty, and staff. Peer interactions can provide a sense of support as well as be important in transferring heuristic knowledge about navigating graduate school (Gardner, 2007; Knight, Hall, & Green-Powell, 2014; Padilla, 1999; Tinto, 1982; Lovitts, 2001). Interactions with faculty (beyond just the advisor) contribute to a student’s sense of belonging in the broader academic community. Further, faculty and staff interactions shape a student’s perception of departmental and institutional support, which can impact retention (Astin, 2014; Bean, 1980; Golde, 2005; Knight et al., 2014; Zhou & Okahana, 2019)

#### Finances

Financial stability and support is also a core factor in student persistence and eventual completion. STEM PhD students are in a unique situation where their tuition is typically paid for them and they receive a stipend or salary to pay for personal expenses such as housing and meals. This funding may come in the form of employment as a teaching assistant (TA) or research assistant (RA) or as a stipend from a fellowship. But even with these sources of support, there can be great disparities in financial stability depending on amount of support and a student’s particular circumstances. Unsurprisingly, the types and amount of funding play a role in doctoral students’ decisions to remain in graduate school (Ampaw & Jaeger, 2012; Herman, 2008; Martinez, Ordu, Sala, & McFarlane, 2013)

#### Preparedness

Graduate study, particularly doctoral research, is a huge undertaking that differs significantly from undergraduate education. Many students enter graduate school without a robust understanding of the expectations thereof and are unaware that the skill set to succeed is very different from that needed in undergraduate studies (Brill, Balcanoff, Land, Gogarty, & Turner, 2014). Students from underrepresented backgrounds may be further disadvantaged with fewer personal connections to graduate students and professors who can share the expectations and “hidden curriculum” of graduate school.

While previous studies have identified these core reasons why graduate students leave, more evidence is needed for student support strategies that can counteract these factors and increase student retention.

### Description of the CE program

CE is a bridge program at UCLA that aims to support URG doctoral students in STEM fields during their transition to graduate studies. Incoming students are nominated by their home department, and then a subset of nominees is selected to participate. The program began in 2008 with just a few students per year. However, since 2010, CE cohorts have ranged from 10 to 44 students, with recent cohorts averaging near 40 students. It should be noted that in 2019, the program expanded to include a few social science students, and in 2020 it also included humanities students. But given the history of the program and the primary constituency, this study focuses solely on STEM students. The program was initially funded by a National Science Foundation (NSF) Alliances for Graduate Education and the Professoriate (AGEP) grant and is currently funded by UCLA’s Division of Graduate Education.

CE students participate in an intensive six-week schedule with activities and programming that address a wide variety of skills and support for graduate students. In 2021, the program was conducted remotely on Zoom due to COVID-19 precautions. Students were expected to participate in the program full-time (40 hours a week); eight of those hours consisted of structured professional development / soft skills programming, and the rest of the time was spent performing research. The structured time included workshops on topics such as managing the advising relationship, writing grant and fellowship applications, and managing mental well-being. Students also participated in a journal club for their discipline that met four times over the course of the program. Additionally, there was a session with a panel of past CE student participants and another panel of faculty who were first-generation students. There was also built-in time for students to get to know each other and ask questions of program leaders. The schedule for the 2021 program can be found in supplemental materials (Table S1) Outside of structured events, students were expected to begin doing research guided by their advisor. The research aspect of the program could be remote, but some students did their research in person. Additionally, students worked with their advisor to write a research proposal by the end of the CE program that could be used to apply for a fellowship such as the NSF Graduate Research Fellowship Program. Additionally, students were funded during the program with a $6,000 stipend.

### Objectives of the CE program

Through the various components, CE seeks to meet several broad learning objectives. Specifically, 1) students will learn how to conduct research, 2) students will learn how to communicate their research with others, 3) students will learn how to work with faculty members, and 4) students will be connected to others at UCLA. We recognized that these four objectives are well-designed to address three of the four primary reasons that students leave doctoral study without a PhD: preparedness, advisor-advisee relationship, and socialization. The unaddressed reason for student attrition is finances. While not a stated goal of the program, CE arguably addresses this area of concern with its stipend and its programming on applying to fellowships and how to budget.

Additionally, every program component has specific learning objectives that aim to support students in these four areas (Table S2).

### Our study

In this work, we use quantitative and qualitative survey data of 55+ first-year STEM doctoral students to analyze the impact of the CE bridge program on matriculated URG students. We consider multiple measures of student skills and experiences that align with both the program’s objectives and known causes of graduate student attrition. Given the alignment between the CE program’s objectives and known causes of graduate student attrition, we expect to see better acclimation and possibly retention outcomes for URG doctoral students who participate in the program. Evaluating the impacts of the CE program can determine the effectiveness of program elements as well as provide a basis for iterative improvements. Additionally, others seeking to create better support for PhD students can learn from any strengths or weaknesses revealed by our analysis of the CE program.

## Materials and Methods

This study was done with an approved UCLA Institutional Review Board protocol #21-000756.

### Survey Question Development

Our survey instruments (<u>Supplemental Material</u>) were developed with feedback from the individuals administering the 2021 Competitive Edge (CE) program to ensure we measured metrics related to the program goals. The CE program administers its own feedback surveys focused on curricula assessment; these surveys have significantly fewer questions and a narrower scope. We looked at the responses of past CE administered surveys to gain insight into what parts of the program our own survey tool should focus on. In 2021, the program still administered their own feedback survey but it was conducted separately from the survey we use here.

Additionally, we surveyed faculty and staff who led program workshops on their individual learning objectives, and we consulted the program director on what the overall program objectives were. The majority of our instruments consist of questions written for this study that directly address those objectives. Furthermore, we used a previously published set of questions addressing the topic of sense of belonging (Hermida 2017). The majority of our questions were framed as Likert scale questions asking respondent agreement with a statement ranging from strongly disagree (1) to strongly agree (5). Statements selected reflected the skills and attitudes we hypothesized that CE would increase. For example, one statement read “I can set expectations with my advisor.” Lastly, we collected demographic data with questions modeled closely after the Grad Student Experience in the Research University survey (SERU Consortium 2021).

### Study Population

All students surveyed were first-year doctoral students in STEM fields at the University of California, Los Angeles (UCLA). Definition of STEM fields for this study included social science disciplines (Table S3). The study population consists of three subgroups referred from here on as 1) Competitive Edge (CE), 2) Non-Competitive Edge Under-Represented Group (NonCE URG), and 3) Non-Competitive Edge Well-Represented (NonCE WR). CE students were those that participated in the 2021 Competitive Edge program. To be eligible for CE, students must be a U.S. citizen, U.S. national, permanent resident, or undocumented student who qualifies for nonresident supplemental tuition exemptions under California law AB 540 (University of California Office of Admissions 2023). Thus we limited our results for our two NonCE cohorts to the same citizenship categories. Additionally CE students must have a background that is underrepresented in graduate education. Students in both NonCE groups did not participate in the program (that year or any other). For NonCE students, URG vs WR students were classified by UCLA’s Graduate Division’s definition of URG students. Functionally, this designation separated White, Asian, and mixed-race White and Asian students as WR and all others as URG. UCLA identified URG students through self-reported information on graduate school applications. We identified URG students based on self-reported information on our survey. However, we note that this definition of URG does not apply to every STEM field. For example, Asian students are underrepresented in Ecology (Kou-Giesbrecht 2020). Nonetheless, given that the aims of the program are to address the needs of disadvantaged students, particularly those defined by the university as belonging to these racial and ethnic/racial categories, we find it prudent to evaluate the strengths of the program under those same categories. We present broad self-reported demographic information on our respondents in table 1.

**Table 1.**
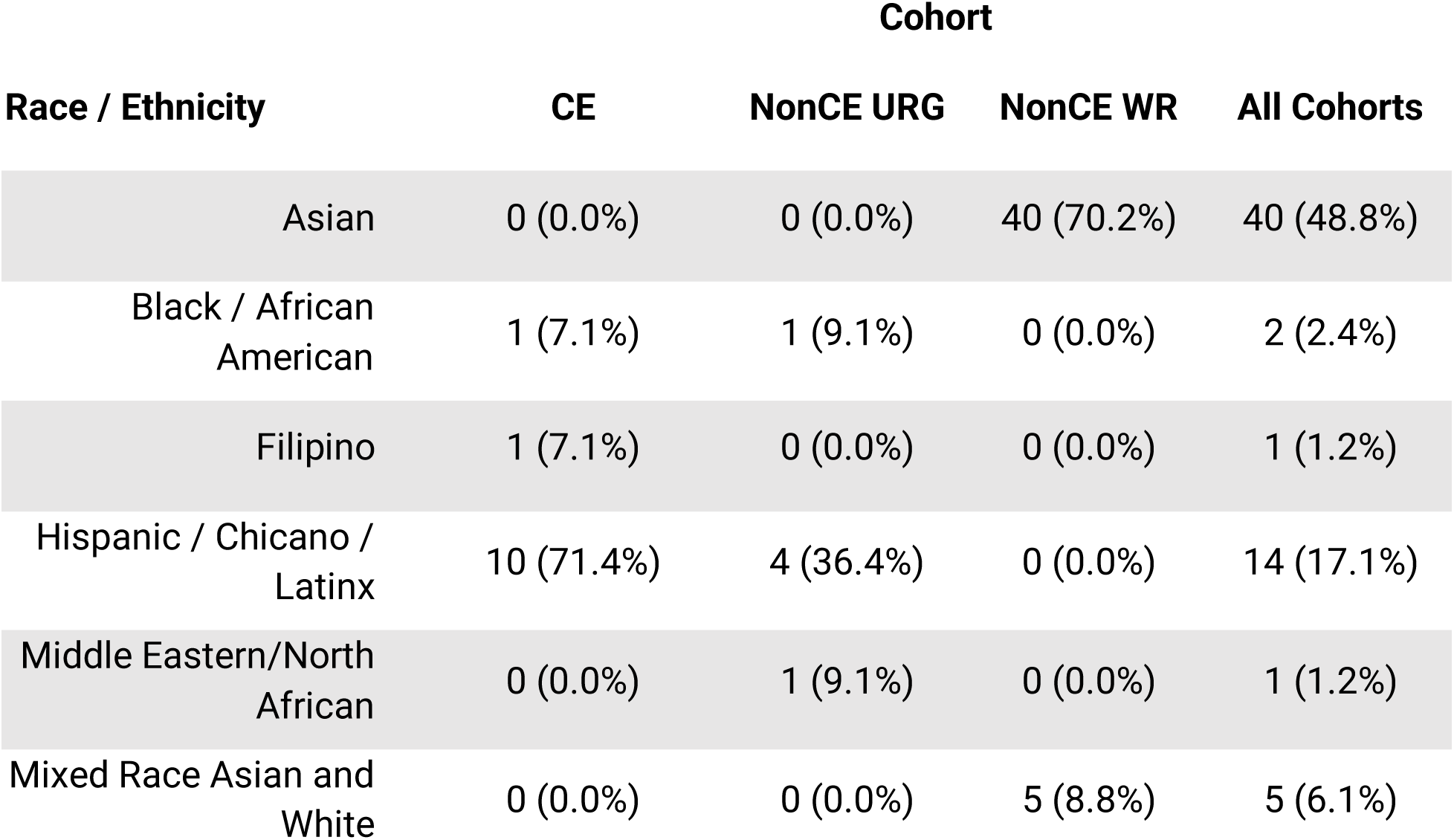
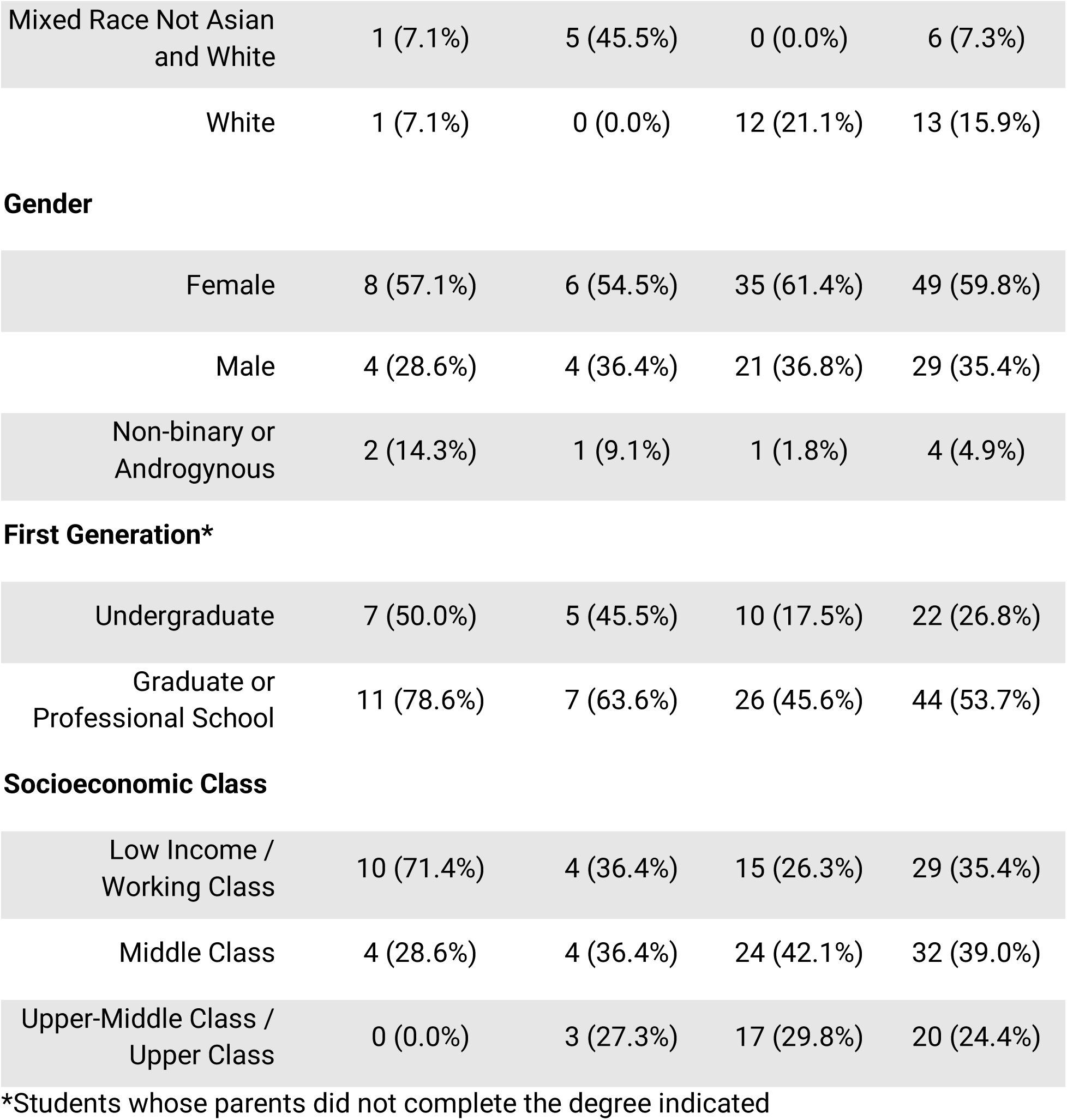
Demographic breakdown of responses by cohort. Numbers represent the total number of responses for a specific cohort and demographic category. Percentages in parentheses show what percentage that number makes up of a given cohort.

| Race / Ethnicity | Cohort |  |  |  |
| --- | --- | --- | --- | --- |
|  | CE | NonCE URG | NonCE WR | All Cohorts |
| Asian | 0 (0.0%) | 0 (0.0%) | 40 (70.2%) | 40 (48.8%) |
| Black / African American | 1 (7.1%) | 1 (9.1%) | 0 (0.0%) | 2 (2.4%) |
| Filipino | 1 (7.1%) | 0 (0.0%) | 0 (0.0%) | 1 (1.2%) |
| Hispanic / Chicano / Latinx | 10 (71.4%) | 4 (36.4%) | 0 (0.0%) | 14 (17.1%) |
| Middle Eastern/North African | 0 (0.0%) | 1 (9.1%) | 0 (0.0%) | 1 (1.2%) |
| Mixed Race Asian and White | 0 (0.0%) | 0 (0.0%) | 5 (8.8%) | 5 (6.1%) |
| Mixed Race Not Asian<br>and White | 1 (7.1%) | 5 (45.5%) | 0 (0.0%) | 6 (7.3%) |
| White | 1 (7.1%) | 0 (0.0%) | 12 (21.1%) | 13 (15.9%) |
| Gender |  |  |  |  |
| Female | 8 (57.1%) | 6 (54.5%) | 35 (61.4%) | 49 (59.8%) |
| Male | 4 (28.6%) | 4 (36.4%) | 21 (36.8%) | 29 (35.4%) |
| Non-binary or<br>Androgynous | 2 (14.3%) | 1 (9.1%) | 1 (1.8%) | 4 (4.9%) |
| First Generation\* |  |  |  |  |
| Undergraduate | 7 (50.0%) | 5 (45.5%) | 10 (17.5%) | 22 (26.8%) |
| Graduate or<br>Professional School | 11 (78.6%) | 7 (63.6%) | 26 (45.6%) | 44 (53.7%) |

|  |  |  |  |  |
| --- | --- | --- | --- | --- |
| Low Income /<br>Working Class | 10 (71.4%) | 4 (36.4%) | 15 (26.3%) | 29 (35.4%) |
| Middle Class | 4 (28.6%) | 4 (36.4%) | 24 (42.1%) | 32 (39.0%) |
| Upper-Middle Class /<br>Upper Class | 0 (0.0%) | 3 (27.3%) | 17 (29.8%) | 20 (24.4%) |

\*Students whose parents did not complete the degree indicated

### Survey Deployment

All surveys were administered via Google forms. Students were incentivized to participate via a raffle for $100 gift cards to Target. In July 2021, we requested the CE program leaders forward our email to the 36 STEM students who made up the 2021 cohort of CE. This was done separately from the CE program’s own feedback surveys. We asked CE students to complete the pre-program questions and had 22 respondents. In June 2022, we again had CE program leaders contact the 2021 cohort students to fill out our end of year-one survey (n = 14).

To survey non-CE students, we had student affairs officers for all STEM PhD programs at UCLA contact current first-year students to take our end of year-one non-CE survey. We also reached out to various graduate student identity-based affinity groups to contact students from racial and ethnic minority backgrounds. According to UCLA STEM Ph.D. program enrollment data, there were 639 first-year PhD students in fall 2021 (UCLA Department of Graduate Education 2023). Of those students, 55% were male, 44% female, and 1% non-binary. Almost a third (31.5%) were international students. UCLA does not release race/ethnicity data for international students. But within domestic students, 20.8% were classified as members of URGs. We received a total of 42 respondents for non-CE students, with eight grouped as from URG backgrounds and 34 grouped from WR backgrounds.

### Survey Response Analysis

To quantitatively analyze our results, we asked CE students to rate how the program improved their experiences across five categories during their first year of doctoral study. The five categories we selected are: **1)** research skills, **2)** sense of belonging, **3)** self confidence, **4)** overall well being, and **5)** interactions with advisors. These categories were hypothesized by us, the CE staff, and contributors to be highly impacted by the program. Furthermore, these categories are highly correlated with success in graduate school (Maher et. al 2020, Holmes et. al 2019, Martinez et. al 2013, Brill et. al 2014).

For additional quantitative insights, we focused on skills relevant to student success that were targeted by the CE program. We asked CE and NonCE students to rate themselves on eight skills: **1)** interacting with faculty, **2)** science communication, **3)** mental wellbeing strategies, **4)** connection to resources, **5)** conducting research, **6)** evaluating journal articles, **7)** financial literacy, and **8)** fellowship application writing. These eight areas were chosen because one or more components of CE focuses on building these skills. For example, students attend a journal club where they learn both how to evaluate literature in their field and how to communicate the findings of those articles. Furthermore, these skills are critical to success in graduate school.

To assess these five experience categories and eight skills, we asked one or more Likert scale questions relating to each one. If multiple questions were incorporated into the score for a specific category or skill, we averaged the answers to all the relevant questions. Additionally, for the eight skills, we calculated students’ growth by subtracting how they rated themselves prior to starting CE or doctoral study (pre-score) from self ratings at the end of their first year of graduate school (post-score).

Our end of year-one surveys concluded with three open ended questions. For the CE students, we asked separately about positive impacts and negative impacts of CE on their doctoral study. We also asked for any additional comments about their first year not addressed by the rest of the survey. For the two NonCE groups, we asked about any programs or experiences that positively impacted and then any that negatively impacted their first year. We also asked for additional comments not addressed elsewhere. These qualitative answers were categorized using an inductive coding method (Thomas 2003). To do this, one author read through all the responses before listing possible categories. They then re-read every response and assigned one or more categories to each response. They consolidated all those themes into 13 broader themes and re-coded all responses with those 13 themes. Those were then collapsed into five final themes: **1)** community support, **2)** financial resources, **3)** mental health, **4)** mentorship/advising, and **5)** skills development (<u>Table S4</u>). The first author then labeled all responses with those classifications. The second author reviewed them and agreed they were consistent with the theme definitions. For the additional comment responses, the text segments were classified as positively or negatively impacting student experiences and combined with the appropriate group of responses. We then tabulated the number of times a given category was present in each cohort’s positive and negative responses.

### Statistical Analysis

Quantitative analysis and plotting of results was carried out in RStudio (RStudio Team 2020). For all statistical tests, an alpha of 0.05 was used. Statistical tests used the T test for comparing differences between two groups. For the statistical comparisons we carried out, we also calculated a power analysis. For the power analysis, we simulated 1,000 data sets per condition for T tests comparing CE vs NonCE URG and CE vs NonCE WR students. We tested effect sizes varying from 0.2 to 2. Effect sizes were defined as the mean of the CE cohort minus the mean of NonCE Cohort then divided by the standard deviation of the NonCE cohort. The means and standard deviations of our simulated data were based on the same parameters from our observed data for NonCE cohorts. To examine the power of each condition, we calculated the proportion of simulations that correctly resulted in a statistically significant T test result of < 0.05.

## Results

### What experiences relating to student success did CE help students with?

Importantly, we asked CE students to rate at the end of their first year how much the program impacted their experiences in 5 key areas. The areas we focused on were **1)** research skills, **2)** sense of belonging, **3)** self-confidence, **4)** overall wellbeing, and **5)** interactions with advisors. Students rated if CE positively influenced their experience with a Likert scale 1 (strongly disagree) to 5 (strongly agree) (Figure 1). We present our areas of focus here in ascending order of mean student response. Research skills had the lowest mean response of 3.7 (between neutral and somewhat agree). CE students predominantly agreed that the program improved their sense of belonging with an average response of 3.9 (between neutral and somewhat agree). All but one student agreed that CE improved their self-confidence with a mean response of 4.2 (somewhat agree). CE students agreed that CE improved their overall wellbeing with a mean response of 4.2 (somewhat agree). CE student respondents unanimously agreed that the program aided their interactions with their advisors (mean = 4.5). CE students reported generally having improved experiences across these five areas relating to student success that they attribute to the CE program. These responses speak to the high value of the CE program.

**Figure 1.**
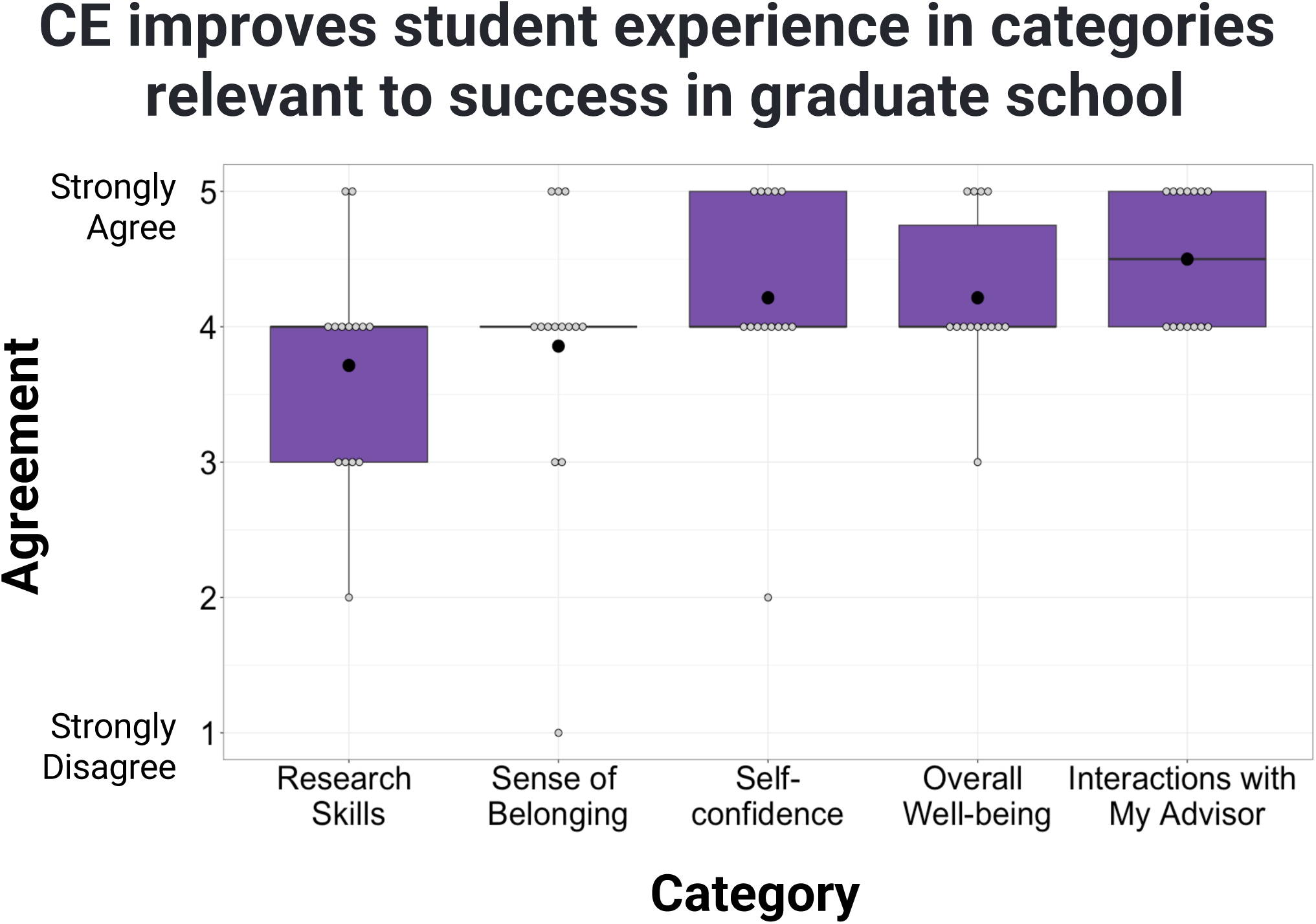
Box plots of CE students’ level of agreement with the statements: “During the 2021-2022 academic year, the Competitive Edge program improved my…” for five categories that the program focuses on improving (x-axis). Responses (y-axis) were on a Likert scale from 1 (Strong Disagree) to 5 (Strongly Agree). Light gray dots represent individual responses while black dots represent mean responses. Thick black lines represent median values.

### What structured components of CE were most helpful?

We asked CE students to select up to three of the most helpful structured components of the program for them. The majority of students reported benefiting from the mental health strategies workshop focused on “Resiliency and Managing Negative Thoughts” (71.4%). They also identified one of the two workshops focused on managing their relationship with their mentor (42.8%). And students did also report benefits from the writing skills workshop on grant and fellowship applications (35.7%).

### CE students reported improvement in skills related to success in graduate school

Students rated themselves before doctoral study and at the end of year-one in eight skills: **1)** interacting with faculty, **2)** science communication, **3)** mental wellbeing strategies, **4)** connection to resources, **5)** conducting research, **6)** evaluating journal articles, **7)** financial literacy, and **8)** fellowship application writing. We calculated students’ change (post score - pre score) in our eight skills for our three cohorts (CE, NonCE URG, and NonCE WR) (Figure 2). CE students reported a larger mean increase in skills than both NonCE cohorts for all skills except financial literacy. For financial literacy, NonCE URG students indicated the most increase in this skill.

**Figure 2.**
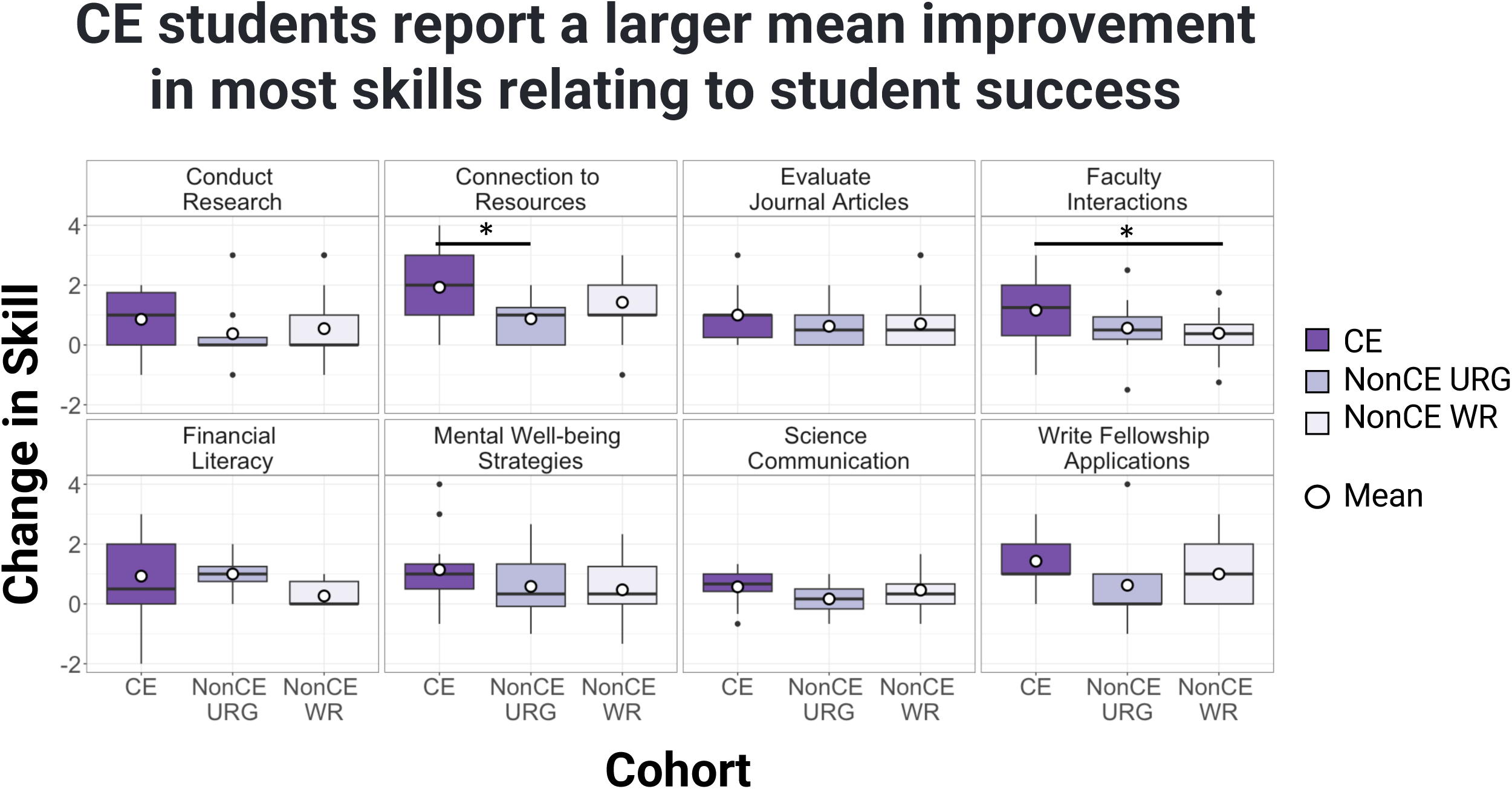
Box plots showing the self-reported change in skills related to success in doctoral studies during the first year of grad school (scale -4 to 4). Cohorts are shown in different colors (CE = dark purple, NonCE URG = purple, and NonCE WR = light purple). Skills we measured are displayed at the top of each panel. White dots represent mean responses and thick black lines represent median values. Black dots represent outliers. Horizontal lines with an asterisk represent T tests that were statistically significant for p-values < 0.05. For all skills but financial literacy, CE students report a higher mean improvement in these skills.

We were most interested in statistically testing if CE students reported more improvements than each of the NonCE cohorts. We did so using a T test to make two pairwise comparisons (CE vs NonCE URG and CE vs NonCE WR) for each skill. We found two statistically significant differences. For CE vs NonCE URG students, CE students reported a larger growth in their connection to resources (T test, p-value = 0.030). For CE vs NonCE WR students, CE students indicated larger growth in working with faculty (T test, p-value = 0.045). All other tests resulted in p-values > 0.05 (Table S5).

Additionally, we used a power analysis to determine the probability that we could accurately detect a difference between our cohorts with our sample sizes plus the observed mean and standard deviation of our control NonCE groups (Figure 1). For comparisons of NonCE students we used the averages of their mean and standard deviation per skill. For URG students the mean was 0.60 and standard deviation was 1.0. WR students had a mean of 0.66 and a standard deviation of 0.80. For our power analysis of comparisons between CE and NonCE URG students, we find that a large effect size > 1.25 would be needed to correctly find a statistical difference in at least 80% of simulated data sets. For CE vs NonCE WR students, a large effect size of 0.75 would be needed to reach 80% accurate statistical tests. Given the means and standard deviations of the NonCE groups, our power analysis suggests we could accurately detect statistical significance in 80% of cases where our CE cohort had a mean response that is > 1.85 units above NonCE URG students or > 1.22 units above NonCE WR students. Our two statically significant cases have a smaller difference in mean than those thresholds, but still may be in the less than 80% of cases where we could detect a true difference.

### What did students say?

Students were asked to identify factors that influenced their first year of graduate school and separate them by those that had positive vs negative impacts. For CE students, there were 9 responses (64.2% of cohort) for positive factors and 5 responses (35.7% of cohort) for negative factors. Among NonCE URG students, 2 responded (18.2% of cohort) with positive and the same 2 (18.2% of cohort) with negative factors. NonCE WR students had 15 responses (26.3% of cohort) for positive and 15 responses (26.3% of cohort) for negative factors.

We coded student responses based on our inductively created categories of **1)** community support, **2)** skills development, **3)** mentorship/advising, **4)** mental health, and **5)** financial resources. A given response could be coded as discussing more than one category. We totaled the number of responses for each category subdivided by cohort and positive vs negative factors (Table 2). The themes are listed above and in the table by order of their popularity ranging from community support with 31 responses to financial resources with 6 responses for all cohorts and positive and negative factors combined.

**Table 2.** Counts of student responses to open ended questions classified by cohort, question, and themes. All students were asked to name factors that contributed positively to their first year of doctoral study. Then they were asked about negative factors. Column titles divide response counts by the three cohorts and/or the question prompt of positive vs negative factors. Five common themes were identified in these responses with inductive coding. These themes are listed in column 1. The final row shows the total number of responses for a given column. A single response could be categorized with discussing multiple themes, so the total responses in a column may be smaller than the total number of times a given topic was discussed by that group.

| Themes | CE |  | NonCE URG |  | NonCE WR |  | All Cohorts |  | Total |
| --- | --- | --- | --- | --- | --- | --- | --- | --- | --- |
|  | Positive | Negative | Positive | Negative | Positive | Negative | Positive | Negative |  |
| Community Support | 8 | 4 | 1 | 1 | 12 | 5 | 21 | 10 | 31 |
| Skills Development | 5 | 2 | 1 | 0 | 4 | 6 | 10 | 8 | 18 |
| Mentorship/Advising | 3 | 0 | 1 | 2 | 3 | 1 | 7 | 3 | 10 |
| Mental Health | 2 | 1 | 0 | 0 | 2 | 4 | 4 | 5 | 9 |
| Financial Resources | 1 | 1 | 0 | 0 | 2 | 2 | 3 | 3 | 6 |
| Response Totals | 9 | 5 | 2 | 2 | 15 | 14 | 26 | 21 | 47 |

The most common theme was community support which appeared in over 65% of responses. All but one CE student who responded to the positive question commented on how CE made them feel supported. One CE student wrote: *“Because of CE, I felt like I belonged at UCLA.”* And 3 of the 4 negative CE responses about community support noted that they think the online delivery of the program hindered their community building within CE. For example: *“…the program being completely virtual made it difficult to connect with other students.”* NonCE students generally referred to student groups and structured university activities that connected them to others at UCLA. One NonCE WR student noted they found community through a student association: *“The engineering graduate student association plans a lot of events which helps us get to know each other and meet new people.”* While another NonCE WR student found connection to be lacking: *“I didn’t feel very connected to my department or the school as a whole…”* Reflecting the importance of community for diversity, a NonCE URG student poignantly wrote, *“I feel like I don’t belong in a lot of the spaces I have been a part of thus far.”*

Skills development was also a popular topic, but referred to both technical and “soft skills” such as achieving work life balance. NonCE WR students in particular commented on skills they wish they had been taught, including interacting with an advisor. For example, “*Interacting with advisors has been a difficult journey for me.*” They also noted programs that were not effective, such as a departmental coding “bootcamp.” CE students also reiterated points about a software workshop that wasn’t particularly relevant to them because their field favors a similar but different software program: “*I would have appreciated an R [software] tutorial day versus a Tableau [software] training.*”

Mentorship and advising proved to be a salient topic for students. CE students noted that it was particularly helpful to begin working with their advisor over the summer. One student wrote, “*I really appreciated the opportunity to do a small research project and start working with my advisor before the beginning of the school year.*” And another added, “*I was also able to publish two manuscripts within my first year because I was able to begin working on them during [CE].*” In contrast, both NonCE URG students and one WR respondent specifically wished for help with navigating their advising relationship. One URG student wrote, “*I need help navigating the relationship with my advisor…*” and the WR student similarly stated, “ *I wish there were more tips and guidance about how to find and interact with an advisor*.” Furthermore, when NonCE WR and URG students cited mentorship as playing a positive role in their first year, they only credited mentorship from sources other than their advisor such as student organizations, senior graduate students, or “younger faculty.”

Mental health was highlighted as impacting first year experiences in nearly 20% of student responses. Two of the 10 CE students focused on how mental health oriented workshops benefited them and the one negative response expressed a desire for more workshops on mental health related topics including “burnout.” In contrast, only two of the 19 NonCE WR students remarked on having support in maintaining their mental health. And four NonCE WR remarked that they were struggling with their mental health. For example, one NonCE WR student wrote: “*It has been very hectic trying to balance coursework with research expectations and finding a project I want to work on.*” and another added, “*I’m super depressed.*”

Lastly, a factor multiple students identified was financial resources. One CE student commented, “*I am very thankful for this program in their financial support…*” But another CE student wrote, “*I was unable to do research in-person [due to constraints with my access to health insurance]. That significantly hindered my ability to make progress in my project.*” While the CE program was conducted online, students had the option to carry out CE research in person. But some mechanism relating to accessing health insurance hindered this student’s experience in the program. For the NonCE cohorts, no URG students mentioned finances. For the four NonCE WR students who mentioned financial resources, three commented specifically on housing. One was grateful for university subsidized family housing. But the other two NonCE WR students commented on the strains of the Los Angeles housing market. For example, “*Looking for affordable housing within a 20 minute walking distance to campus has been very difficult…[Commuting] while living not within walking distance to campus makes it hard to get enough rest and maintain well-being.*”

## Discussion

The research described here highlights ways a diversity-oriented STEM summer bridge program can benefit students during their first year of doctoral study. In particular, quantitative and qualitative analyses of responses from CE students and control cohorts identified data trends supporting a positive impact of CE on research skills and psychosocial traits important for success in graduate school. From the perspective of diversifying STEM graduate programs, we present the benefits reported by CE students in the context of four major causes of graduate student attrition.

### Advisor-Advisee Relationship / Working with Faculty

A positive effect of CE on students’ advisor-advisee relationships was evident throughout our analyses. Notably, CE students unanimously agreed that the program improved their interactions with their advisor (Fig 1). When asking students to rate their skill of interacting with faculty, CE students reported a higher mean growth in working with faculty compared to NonCE URG and WR students (Fig 2). The positive impact was likely driven by two workshops which focused entirely on how to manage interactions with advisors. CE students identified the “Mentoring Up” workshop as the second most helpful CE program component.

Additionally, CE students repeatedly commented on the benefit of starting research training with their advisor over the summer. Summer may be a particularly advantageous time to begin working with an advisor because faculty generally have more availability due to limited teaching and service requirements. While many faculty may take vacation during the summer, and in some disciplines may conduct field work over this time, the CE program requires students to have a faculty member who commits to mentoring the student during the program. So the students are paired with advisors who are explicitly available during the six week program. Thus, compared to NonCE cohorts, CE students in our study had additional training in how to manage their advising relationship and additional time to navigate the interaction.

It is worth noting that multiple NonCE students commented on difficulties interacting with their advisors. One URG and one WR student explicitly wished for more support in navigating this relationship. These observations emphasize the need for this type of support for STEM PhD students and highlight the significance of our finding that all CE respondents viewed the program as specifically assisting them in interacting with their advisors.

### Socialization / Connection to others at UCLA

Receiving social support may be particularly salient to CE students because of their minoritized racial/ethnic identities as well as first-generation status. All but one CE student identified with a URG racial or ethnic identity (Table 2). There is strong evidence that it is important for mentees from URG backgrounds to have mentors with cultural awareness (Thomas 2001, Osula & Irvin 2009, Womack *et al*. 2020). Additionally, 50% of CE students reported being first-generation with regards to college degrees and 78.6% with regards to advanced degrees. In contrast, 17.5% of NonCE WR students were first-generation college students and 45.6% first-generation for post-college education. The lack of familial experience in advanced degree studies may translate to fewer avenues for first-generation students to learn how to navigate the non-technical parts of graduate school. Additionally, we view socialization as linked to mental health. Social connections can offer emotional support, companionship, and a sense of belonging, which can help individuals cope with stress, reduce feelings of loneliness, and enhance overall well-being. Notably, multiple aspects of our analyses show that CE students benefited from the social aspects of CE.

Most CE students agreed that the program improved their sense of belonging, their self-confidence and their overall well-being during their first year (Fig 1). These findings are also reflected in CE students reporting a larger increase in their mental wellbeing strategies and their connection to resources compared to both NonCE groups (Fig 2). When asked for the most helpful components of CE, students’ top answer was a workshop that highlighted social strategies to build self-efficacy (belief that one can do what is necessary to achieve their goals). These results all support a conclusion that CE student socialization was aided by the program.

In all cohorts’ short answer responses, community support was the most common theme. CE students explicitly commented on how the program made them feel more connected at UCLA. While many NonCE URG and WR students described community support they had found, some described a lack of belonging. Relatedly, students from the CE and NonCE WR cohorts commented on mental health topics. CE students reaffirmed how the self-efficacy workshop helped them as well as a workshop on resilience and negative thoughts. Two NonCE WR students remarked on the support they found for their mental health, but four described ways they needed more support.

The very nature of the CE program gave this cohort of students access to URG peers and invested faculty and staff, which created a natural space for these students to feel connected and supported from the start of their program. Also, the workshops the CE students highlighted focused on non-technical or so-called “soft skills.” These types of skills are not usually taught explicitly in graduate education. Instead, most traditional elements of graduate education, such as coursework and advising from PIs, is focused on developing the critical technical skills needed for the research field. Notably, CE student open-ended responses specifically commented that the program’s focus beyond technical skills was a significant benefit of the program. This speaks to students’ desire and need to develop skills that contribute to graduate student success beyond knowledge of their field of study. Thus it is not surprising that CE students valued support in areas that are not typically a focus of STEM doctoral study.

### Finances

Finances may be particularly relevant to the >70% CE students who self-reported being from low-income and working class socioeconomic backgrounds. This contrasts with ∼36% of NonCE URG students and ∼26% of NonCE WR students with these backgrounds. For the duration of the program, CE students are paid a $6,000 stipend for living expenses and possibly to offset some of their moving costs. For reference, UCLA graduate student teaching assistants in Fall 2023 would be paid a minimum of only $3,608 for the same time period (UAW Local 2865 2023). The pay above the university minimum that CE provides may be a significant boost to financial stability for CE students during the critical period of transition to their doctoral study. Such an effect was noted by one CE student’s open-ended response stating gratitude for the program’s financial support. Additionally, CE students are encouraged to move to Los Angeles to conduct their research in person. As the program occurs in the summer, this may give CE students greater potential to find housing at a time when there is less competition from incoming students for housing, including UCLA subsidized housing, near campus. Lastly, CE students have the potential for better financial support in the future from a boost in writing skills from the CE workshop on fellowship applications (discussed in more detail under preparedness).

### Preparedness / Doing and communicating research

In addition to the topics described above, students need to know and develop skills that are important for success in graduate school. We find evidence in multiple of our analyses that CE students developed important skills, including doing and communicating research.

We asked CE students if they thought that the program improved their research skills. The mean response of students was between neutral and somewhat agree (Fig 1). This moderate response likely reflects the limited opportunity for students to delve in depth into research in a program of only 6 weeks. Even so, the most common response was “somewhat agree” (4) suggesting that many students believe their overall research skills benefited from the program.

When asked about the most helpful program components, CE students identified the grant and fellowship writing workshop as one of the top three. Better proficiency in writing can help students more easily secure funding for themselves through fellowships and their research through grants. Additionally, many doctoral programs also have milestones such as a dissertation proposal which has similarities to grant applications. Being able to effectively communicate one’s research plans will likely help these students meet these milestones more easily. Lastly, the final milestone in doctoral study is of course completion of the dissertation itself. The boost in writing skills that CE students gain during their first year has the potential for large long-term advantages.

Lastly, we found a striking trend that CE students reported a larger mean improvement in seven of the eight skills relating to student success compared to NonCE students (Fig 2). The eight skills we focused on were chosen both because the CE program specifically seeks to support students in these areas and because the skills are important for one or more metrics of doctoral student success. Specifically we found statistically significant differences in connection to resources relative to NonCE URG students and interactions with faculty relative to NonCE WR students. However, we note that our statistical power was limited (Fig S1) likely due to our small sample sizes. So we argue that the overall trend of CE students indicating more growth in almost all skills we measured is worth consideration.

### Limitations and future directions

Small sample size limits this study, especially statistical inferences. The number of CE respondents was 14 students, or about one-third of eligible CE students. We chose to focus on a single cohort of CE students because the leadership, instructors, and program topics have varied between years. These variations introduce confounding differences between cohorts that would complicate analysis of data from larger sample sizes obtained by surveying multiple cohorts. The 11 NonCE URG student respondents was also a small sample. This reflects the inherently smaller proportion of the overall student body that URG students comprise by definition. This was also a lower response rate than CE students, likely because NonCE students have no invested interest in the program.

It is worth noting that students are not randomly selected for the CE program. They must be nominated by their department and then selected by a committee. Because of the non-random nature of selecting students to participate in CE there are some uncontrolled variations between our study groups.

We also acknowledge that in our skills comparisons between CE and NonCE groups, there are many paths for students to acquire and develop the skills we studied. While the CE program directly targets these skills and the quantitative data show a trend of more CE students improving than NonCE URG students, CE is not the only way that gap could have developed. However, because the trend occurs across all but one skill, the parsimonious interpretation is that CE conferred an advantage to participating students over their NonCE peers.

In order to explore impacts of CE further, we suggest future studies look across multiple cohorts of CE students. With sufficient sample size, the confounding differences of program and environmental differences could be more easily controlled for in statistical analyses. For example, an ordered logistic regression could incorporate program year as a variable in the statistical model. Additionally, other demographic information such as gender and first generation status could be tested as model components as well.

Lastly, a longitudinal approach should incorporate long term metrics of success for CE students. A particular metric of interest would be attrition, given the alignment between CE’s goals and the causes of graduate student attrition. Future studies could also track other outcomes such as publication record and time to degree. While the primary focus of CE is to assist students during their acclimation to graduate school, it would be prudent to explore the potential long-term effects of beginning doctoral study with a “competitive edge”.

## Conclusion

The Competitive Edge (CE) program at the University of California, Los Angeles (UCLA) seeks to support first year doctoral students from historically excluded and underrepresented groups (URGs). Survey results of CE and NonCE first year PhD students at UCLA suggest that CE achieved the goal of better preparing program participants. Specifically, we found ways that the CE program addressed four major causes of graduate student attrition: 1) advisor-advisee relationship, 2) socialization, 3) finances, and 4) preparedness. *Advising Relationship:* A higher percentage of CE students than their nonCE peers reported improvements in managing interactions with their advisor. Furthermore, all CE respondents indicated that the program specifically helped those interactions. *Socialization:* The majority of CE students agreed that the program improved their sense of belonging and overall well-being. *Finances:* CE students received a significant stipend and were grateful for that financial support. They also reported benefiting greatly from a workshop on fellowship writing, which could improve future funding prospects. *Preparedness:* In seven of eight key skills for graduate students, proportionally more CE students improved compared to their NonCE peers. Based on the program’s impact in these areas, we anticipate the program having long term effects on participants’ retention and success in graduate school, a hypothesis that warrants longitudinal studies of multiple CE student cohorts. Given the positive results of the CE program at UCLA, the program’s model could be used to build or improve upon institutional support for doctoral students from URGs at other institutions.

## Acknowledgements

The authors thank the following individuals for their valuable contributions to this research project. Beverly Yanuaria and Andrew Rameriz for their assistance in collecting program objectives, survey design, and communication to CE students. Dr. Rhiannon Little-Surowski for aiding in survey design and communication to students. Dr. Jaana Juvonen and Stephanie Dolbier for feedback on survey design and data analysis. Dr. Casey Shapiro for substantial feedback on survey design, data analysis, and manuscript writing. Dr. Greg Payne and Dr. Lawrence Evalyn for valuable feedback on our manuscript. Dr. Gabe Hassler for guidance with statistical analyses. We would like to acknowledge the support of the UCLA Division of Graduate Education. This material is based upon work supported by the National Science Foundation Graduate Research Fellowship Program under Grant No. DGE-2034835 (awarded to Author Christina Del Carpio). Any opinions, findings, and conclusions or recommendations expressed in this material are those of the authors and do not necessarily reflect the views of the National Science Foundation. Lastly, the authors acknowledge our presence on the traditional, ancestral, and unceded territory of the Gabrielino/Tongva peoples.

## References

Ampaw, F. D., & Jaeger, A. J. (2012). Completing the Three Stages of Doctoral Education: An Event History Analysis. Research in Higher Education, 53(6), 640– 660. 10.1007/s11162-011-9250-3

Astin, A. W. (2014). Student involvement: A developmental theory for higher education. College Student Development and Academic Life: Psychological, Intellectual, Social and Moral Issues, (July), 251–263.

Bean, J. P. (1980). Dropouts and turnover: The synthesis and test of a causal model of student attrition. Research in Higher Education, 12(2), 155–187. 10.1007/BF00976194

Bowlin, L., Sweat, K., Watts, S., & Throne, R. (2017). Agency, Socialization, and Support: A Critical Review of Doctoral Student Attrition. Online Submission.

Brill, J. L., Balcanoff, K. K., Land, D., Gogarty, M., & Turner, F. (2014). Best Practices in Doctoral Retention: Mentoring. Higher Learning Research Communications, 4(2), 26. 10.18870/hlrc.v4i2.186

Devine, K., & Hunter, K. (2016). Doctoral Students’ Emotional Exhaustion and Intentions to Leave Academia. International Journal of Doctoral Studies, 11, 035–061. 10.28945/3396

Gardner, S. K. (2007). “I heard it through the grapevine”: Doctoral student socialization in chemistry and history. Higher Education, 54(5), 723–740. 10.1007/s10734-006-9020-x

Girves, J. E., & Wemmerus, V. (1988). Developing Models of Graduate Student Degree Progress. The Journal of Higher Education, 59(2), 163–189. 10.1080/00221546.1988.11778320

Golde, C. M. (2005). The role of the department and discipline in doctoral student attrition: Lessons from four departments. Journal of Higher Education, 76(6), 669–700. 10.1080/00221546.2005.11772304

Hardré, P. L., Liao, L., Dorri, Y., & Beeson Stoesz, M. A. (2019). Modeling American graduate students’ perceptions predicting dropout intentions. International Journal of Doctoral Studies, 14, 105–132. 10.28945/4161

Herman, C. (2008). Obstacles to success – doctoral student attrition in South Africa Research on doctoral attrition. Africa, 40–52.

Hermida, A. (2017). Everyday Oppression: The Challenges of Belonging for Underrepresented Doctoral Students at a Predominantly White Institution. [Doctoral dissertation, University of Minnesota]. Core. https://core.ac.uk/display/211352927?utm_source=pdf&utm_medium=banner&utm_campaign=pdf-decoration-v1

Knight, L., Hall, T., & Green-Powell, P. (2014). An Analysis of Historically Black Colleges and Universities Student Retention and Attrition Efforts, 1(8), 123–138.

Kou-Giesbrecht, Sian. “Asian Americans: the forgotten minority in ecology.” The Bulletin of the Ecological Society of America 101.3 (2020): e01696.

Lovitts, B. E. (2001). Leaving the Ivory Tower: The Causes and Consequences of Departure from Doctoral Study. Lanham, Maryland: Rowman & Littlefield Publishers, Inc.

National Academies of Sciences, Engineering, and Medicine. 2019. The Science of Effective Mentorship in STEMM. Washington, DC: The National Academies Press. 10.17226/25568.

Maher, M. A., Wofford, A. M., Roksa, J., & Feldon, D. F. (2017). Exploring Early Exits: Doctoral Attrition in the Biomedical Sciences. Journal of College Student Retention: Research, Theory & Practice, 152102511773687. 10.1177/1521025117736871

Martinez, E., Ordu, C., Sala, M. R. D., & McFarlane, A. (2013). Striving to obtain a school-work-life balance: The full-time doctoral student. International Journal of Doctoral Studies, 8, 39–59. 10.28945/1765

Mccoy, D. L., Winkle-wagner, R., & Winkle-wagner, D. L. M. R. (2020). Bridging the Divide: Developing a Scholarly Habitus for Aspiring Graduate Students Through Summer Bridge Programs Participation Bridging the Divide: Developing a Scholarly Habitus for Aspiring Graduate Students Through Summer Bridge Programs Participation, 56(5), 423–439.

Meara, K. O., Griffin, K. A., & Robinson, T. (2017). Sense of Belonging and Its Contributing Factors in Graduate Education. International Journal of Doctoral Studies, 12, 251–279. 10.28945/3903

Osula, B., & Irvin, S. M. (2009). Cultural awareness in intercultural mentoring: A model for enhancing mentoring relationships. International Journal of Leadership Studies, 5(1), 37–50.

Padilla, R. V. (1999). College Student Retention: Focus on Success. Journal of College Student Retention: Research, Theory & Practice, 1(2), 131–145. 10.2190/6w96-528b-n1kp-h17n

Peteet, B., Bridge, E., Ethnic, P., Ronald, T., Postbaccalaureate, E. M., Program, A., Preparation, I. (2016). How graduate school bridge programs can help increase diversity in STEM subject admission, 1–4.

Rockinson-Szapkiw, A. J. (2019). Toward understanding factors salient to doctoral students’ persistence: The development and preliminary validation of the doctoral academic-family integration inventory. International Journal of Doctoral Studies, 14, 237–258. 10.28945/4248

RStudio Team (2020). RStudio: Integrated Development for R. RStudio, PBC, Boston, MA URL http://www.rstudio.com/.

Ruud, C. M., Saclarides, E. S., George-Jackson, C. E., & Lubienski, S. T. (2018). Tipping Points: Doctoral Students and Consideration of Departure. Journal of College Student Retention: Research, Theory and Practice, 20(3), 286–307. 10.1177/1521025116666082

SERU Consortium. gradSERU Survey Design. (2021). Retrieved March 8, 2021, from https://cshe.berkeley.edu/seru/about-seru/seru-surveys/gradseru-survey-design

Sowell, R., Allum, J., & Okahana, H. (2015). Doctoral Initiative on Minority Attrition and Completion. 10.1145/1401890.1402023

Sowell, R., Zhang, T., Redd, K., & King, M. F. (2008). Analysis of baseline program data from the Ph.D. completion project.

Thomas, D. A. (2001). Race matters. Harvard Business Review. April.

Tinto, V. (1982). Limits of Theory and Practice in Student Attrition. The Journal of Higher Education, 53(6), 687–700. 10.1080/00221546.1982.11780504

UAW Local 2865. (2023). Academic Student Employees (ASE) Contract. Retrieved May 26, 2023 from https://uaw2865.org/ase-contract/

UCLA Department of Graduate Education. (2023). Retrieved May 20, 2023. https://go.grad.ucla.edu/

University of California, Office of Undergraduate Admissions. (2023). AB 540 Nonresident Tuition Exemption. Retrieved June 5, 2023, from https://admission.universityofcalifornia.edu/tuition-financial-aid/tuition-cost-of-attendance/ab-540-nonresident-tuition-exemption.html

Weiss, C. S. (1981). The Development of Professional Role Commitment Among Graduate Students. Human Relations, 34(1), 13–31.

Womack, V. Y., Wood, C. V., House, S. C., Quinn, S. C., Thomas, S. B., McGee, R., & Byars-Winston, A. (2020). Culturally aware mentorship: Lasting impacts of a novel intervention on academic administrators and faculty. PloS one, 15(8), e0236983.

Zhou, E., & Okahana, H. (2019). The Role of Department Supports on Doctoral Completion and Time-to-Degree. Journal of College Student Retention: Research, Theory and Practice, 20(4), 511–529. 10.1177/1521025116682036

