## Supplemental Materials for "UCLA’s Competitive Edge Program Provides an Advantage to STEM Doctoral Students from Historically Excluded and Underrepresented Groups"

**Supplemental Material**

*Supplemental Text*

Surveys used can be found at the following links.

*CE Pre-Program Survey:*
https://drive.google.com/file/d/1Q8F-XHAp9TgcW664yHeOJ67KxMlo0p5m/view?usp=share_link

*CE Year-End Survey:*
https://drive.google.com/file/d/1-q7_IE9ScneFwNyhgQgOapHBp62YhUVj/view?usp=share_link

*NonCE Year-End Survey:*
https://drive.google.com/file/d/1ggTUCp0HBzLkwm3StUQlqkRIW-mWNjM2/view?usp=share_link

| Day | Time | Topic |
|---|---|---|
| | **Week 1** | |
| Mon | 10am - 12pm | Welcome/Orientation |
| Tue | 10am - 12pm | Mentoring Part 1: Mentoring Up |
| Wed | 10am - 12pm | Homeroom* |
| Th | 10am - 12pm | Introduction to Campus and Resources |
| | **Week 2** | |
| Mon | 10am - 12pm | Literature Searching and Reference Management |
| Tue | 10am - 12pm | Writing Successful Grant and Fellowship Applications |
| Wed | 10am - 12pm | Homeroom |

| | | |
|---|---|---|
| Th | 10am - 11am | CE Alumni Panel |
| Th | 11am-12pm | Homeroom |
| | **Week 3** | |
| Mon | 10am - 12pm | Introduction to Tableau |
| Tue | 10am - 12pm | Journal Club Orientation |
| Wed | 10am - 12pm | Resiliency/Negative Cognitive Thoughts |
| Th | 10am - 11am | Mentoring Part II: Effective Communication and Managing Expectations |
| Th | 11am-12pm | Homeroom |
| | **Week 4** | |
| Tue | 10am - 11am | First Generation Faculty Panel |
| Tue | 11am-12pm | Homeroom |
| Wed | 10am - 12pm | Journal Club |
| Th | 10am - 11am | Self-Efficacy |
| Th | 11am-12pm | Homeroom |
| | **Week 5** | |
| Tue | 10am - 11am | Grad School on a Budget |
| Tue | 11am-12pm | Fellowships and Extramural Funding |
| Wed | 10am - 12pm | Journal Club |
| Th | 10am - 12pm | Homeroom |
| | **Week 6** | |
| Tue | 10am - 12pm | Homeroom |
| Wed | 10am - 12pm | Journal Club |
| Th | 10am - 12pm | Program Closing |

**Table S1.** 2021 CE Program schedule. *Homeroom was informal sessions for the CE cohort to discuss academic and non-academic topics. During homeroom, students were typically given a prompt to begin discussion.

| Program Component | Learning Objectives | Area(s) of Student Attrition Addressed |
|---|---|---|
| CE Alumni Panel | Develop a relationship with members of other CE cohorts<br><br>Learn strategies and tips to be a successful graduate student | Preparedness<br>Socialization |
| First-Generation Faculty Panel | Learn strategies and tips to be a successful graduate student<br><br>See representation of different identities among faculty | Preparedness<br>Socialization |
| Graduate School on a Budget | Learn how to budget a typical UCLA student stipend/income | Finances |
| Homeroom | Develop a connection to CE cohort | Socialization |
| Introduction to Campus Resources | Learn about resources available to students at UCLA | Socialization |
| Introduction to Tableau | Become familiar with Tableau software<br><br>Learn how to do fundamental processes in Tableau | Preparedness |

| | | |
|---|---|---|
| Journal Club | Identify a high impact primary research paper in the field that will be of general interest to researchers in life, biomedical and social sciences.<br><br>Present a clear, concise, and critical oral presentation of a high impact research paper in the field that is understandable to a general audience of scientists.<br><br>Prepare slides that are well-organized, uncluttered, legible, and focused on a single important concept or finding.<br><br>Use an oral presentation style that engages the audience.<br><br>Critically evaluate the strengths and weaknesses of research described in a primary research paper.<br><br>Provide constructive feedback to peers on their presentation skills. | Preparedness |
| Literature Searching and Reference Management | Recall how to contact the library, locate and access relevant literature in a research database<br><br>Identify Zotero as a way to save, organize, and cite research literature | Preparedness |
| Mentoring Workshops | Developing strategies for the first interaction(s)<br><br>Working to set a tone for the relationship<br><br>Learning strategies for managing expectations | Advisor-Advisee Relationship |
| Resiliency/Negative Cognitive Thoughts | Learn about the factors that influence resilience such as cognitive distortions with a focus on imposter syndrome.<br><br>Learn how to manage cognitive distortions and how to foster coping strategies that build resilience during times of crisis. | Socialization |

| | | |
|---|---|---|
| Self-Efficacy | Define Self-efficacy<br><br>Learn about the four sources of self-efficacy and how to utilize them to bolster your own self-efficacy. | Socialization |
| Writing Successful Grant and Fellowship Applications | Understand effective writing strategies for fellowship essays (personal statements and research proposals)<br><br>To understand the different funding opportunities students are eligible for and how the funding packages work. | Finances<br>Preparedness |

**Table S2.** Structured CE program components are listed with their learning objectives and areas of graduate student attrition that they address. For program components previously offered, learning objectives were reported by the person who previously led that component. Learning objectives for new components were written by the authors in consultation with CE program leadership.

| | |
|---|---|
| Anthropology | Geography |
| Archaeology | Geophysics & Space Physics |
| Astronomy and Astrophysics | Health Policy & Management |
| Atmospheric and Oceanic Sciences | Immunity, Microbes & Molecular Pathogenesis |
| Biochemistry, Biophysics & Structural Biology | Information Studies |
| Bioengineering | Materials Science & Engineering |
| Bioinformatics | Mathematics |
| Biomathematics | Mechanical and Aerospace Engineering |
| Biostatistics | Medical Informatics |
| Cell & Developmental Biology | Molecular Pharmacology |
| Chemical Engineering | Molecular, Cellular & Integrative Physiology |
| Chemistry | Neuroscience |
| Civil Engineering | Neuroscience |

| | |
|---|---|
| Community Health Sciences | Nursing |
| Computer Science | Oral Biology |
| Ecology and Evolutionary Biology | Physics |
| Economics | Physics & Biology in Medicine |
| Electrical & Computer Engineering | Planetary Science |
| Environment and Sustainability | Political Science |
| Environmental Health Sciences | Psychology |
| Epidemiology | Public Health |
| Gene Regulation, Epigenomics & Transcriptomics | Sociology |
| Genetics & Genomics | Statistics |
| Geochemistry | Urban Planning |

**Table S3.** List of the PhD programs at UCLA that were categorized as STEM fields.

| Themes | Description | Examples |
|---|---|---|
| **Community Support** | Receiving support from a community member. Feeling a sense of belonging to a community. Campus resources are included here. Many levels of community exist such as faculty, students, student groups, departmens, etc. This does not include financial support. | "Because of CE, I felt like I belonged at UCLA."<br><br>"My first year is graduate school has been difficult and it's not because I don't have the skills to and tools to navigate grad school, but rather because my program does not adequately support me or other minoritized students." |
| **Financial Resources** | Having financial resources can be personal (eg stipend or fellowship) or for research (eg grants). Resources influenced by finances include housing, commute, and child care. | "Looking for affordable housing within a 20 minute walking distance to campus has been very difficult."<br><br>"I was unable to do research in- |

| | | |
|---|---|---|
| | | person since I didn't have California health insurance" |
| **Mental Health** | Topics relating to mental health such as imposter syndrome, anxiety, depression, and self-efficacy. Work life balance is also included. | "It has been very hectic trying to balance coursework with research expectations and finding a project I want to work on"<br><br>"I'm super depressed." |
| **Mentorship/ Advising** | Mentorship can be from any community member, but many responses are focused on mentorship from an advisor/PI. | "Interacting with advisors has been a difficult journey for me. "<br><br>"I wish there were more tips and guidance about how to find and interact with an advisor." |
| **Skills Development** | Skills development includes fostering technical ("hard" skills) and "soft" skills. The modes of learning or improving skills most mentioned include workshops, journal clubs, and courses. | "[CE] did not solely focus on preparing us to conduct research but focused on other aspects that are necessary yet forgotten such as building self-efficacy and setting expectations with a new mentor."<br><br>"I wish there'll be more opportunities for workshop that can be helpful to my research." |

**Table S4.** List of the five major themes observed through deductive coding of open - ended response questions. Definitions and examples of each theme are provided.

| Skill | Cohort | Mean | Min | Max | Standard Deviation | p-value for T test vs CE |
|---|---|---|---|---|---|---|
| Conduct research | CE | 0.9 | -1.0 | 2.0 | 0.9 | - |
| | NonCE URG | 0.4 | -1.0 | 3.0 | 1.2 | 0.34 |
| | NonCE WR | 0.5 | -1.0 | 3.0 | 0.9 | 0.31 |
| Connection to resources | CE | 1.9 | 0.0 | 4.0 | 1.3 | - |
| | NonCE URG | 0.9 | 0.0 | 2.0 | 0.8 | 0.030 |
| | NonCE WR | 1.4 | -1.0 | 3.0 | 1.1 | 0.21 |
| Evaluate journal articles | CE | 1.0 | 0.0 | 3.0 | 0.9 | - |
| | NonCE URG | 0.6 | 0.0 | 2.0 | 0.7 | 0.30 |
| | NonCE WR | 0.7 | 0.0 | 3.0 | 0.9 | 0.30 |
| Faculty interactions | CE | 1.2 | -1.0 | 3.0 | 1.3 | - |
| | NonCE URG | 0.6 | -1.5 | 2.5 | 1.2 | 0.28 |
| | NonCE WR | 0.4 | -1.3 | 1.8 | 0.6 | 0.045 |
| Financial literacy | CE | 0.9 | -2.0 | 3.0 | 1.5 | - |
| | NonCE URG | 1.0 | 0.0 | 2.0 | 0.8 | 0.88 |
| | NonCE WR | 0.3 | 0.0 | 1.0 | 0.4 | 0.12 |
| Mental wellbeing strategies | CE | 1.1 | -0.7 | 4.0 | 1.2 | - |
| | NonCE URG | 0.6 | -1.0 | 2.7 | 1.2 | 0.30 |
| | NonCE WR | 0.5 | -1.3 | 2.3 | 0.9 | 0.07 |
| Science communication | CE | 0.6 | -0.7 | 1.3 | 0.6 | - |

| | | | | | | |
|---|---|---|---|---|---|---|
| | NonCE URG | 0.2 | -0.7 | 1.0 | 0.6 | 0.16 |
| | NonCE WR | 0.5 | -0.7 | 1.7 | 0.5 | 0.53 |
| Writing fellowship applications | CE | 1.4 | 0.0 | 3.0 | 0.8 | - |
| | NonCE URG | 0.6 | -1.0 | 4.0 | 1.5 | 0.19 |
| | NonCE WR | 1.0 | 0.0 | 3.0 | 0.9 | 0.11 |

**Table S5.** Table of parameters for student self-reported changes in eight skills relating to success in graduate school.

### Power analysis shows large effect size needed to find statistical significance

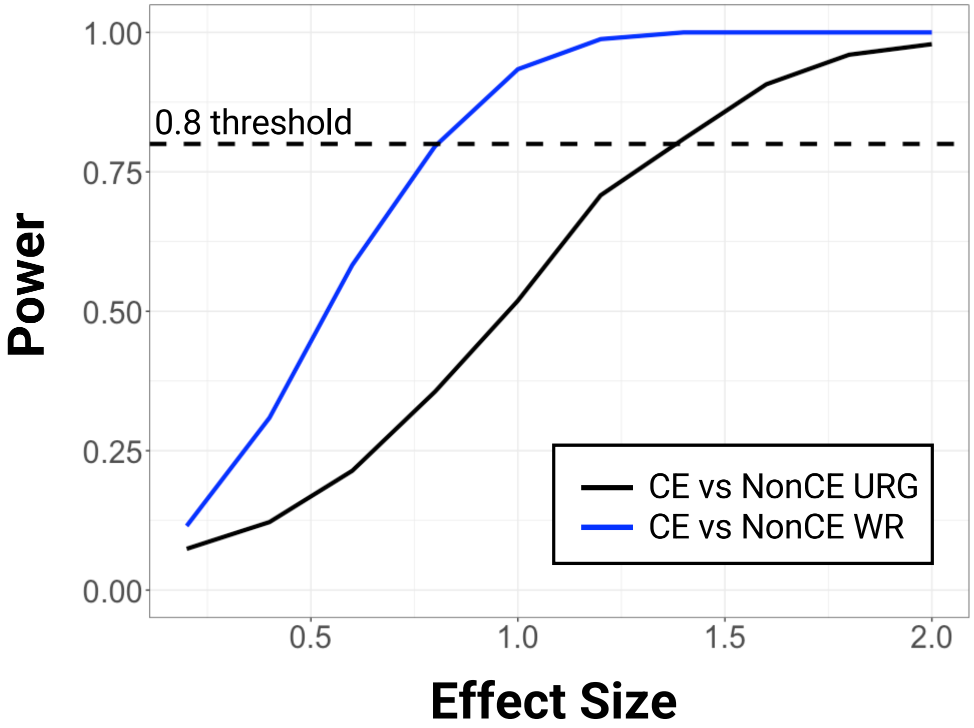


**Figure S1.** Plot of the power for a T test on simulated datasets with varying effect sizes for comparing CE students responses to NonCE URG (black solid line) and NonCE WR (blue solid line) responses. For each effect size tested, we simulated 1,000 data sets. Each data set simulated had a sample size, mean, and standard deviation based on the observed parameters of our collected data. The dashed line at a power threshold of 0.8 shows the effect size needed to accurately detect a difference between CE students' responses and NonCE students' responses is large: >0.75 for NonCE WR and >1.25 for NonCE URG students.
